# Catalytic properties of RNA polymerases IV and V: accuracy, nucleotide incorporation and rNTP/dNTP discrimination

**DOI:** 10.1101/117184

**Authors:** Michelle Marasco, Weiyi Li, Michael Lynch, Craig S. Pikaard

## Abstract

Catalytic subunits of DNA-dependent RNA polymerases of bacteria, archaea and eukaryotes share hundreds of ultra-conserved amino acids. Remarkably, the plant-specific RNA silencing enzymes, Pol IV and Pol V differ from Pols I, II and III at ~140 of these positions, yet remain capable of RNA synthesis. Whether these amino acid changes in Pols IV and V alter their catalytic properties in comparison to Pol II, from which they evolved, is unknown. Here, we show that Pols IV and V differ from one another, and Pol II, in nucleotide incorporation rate, transcriptional accuracy and the ability to discriminate between ribonucleotides and deoxyribonucleotides. Pol IV transcription is notably error-prone, which may be tolerable, or even beneficial, for biosynthesis of siRNAs targeting transposon families *in trans.* By contrast, Pol V exhibits high fidelity transcription, suggesting a need for Pol V transcripts to faithfully reflect the DNA sequence of target loci in order to recruit siRNA-Argonaute protein silencing complexes.

## Introduction

In all eukaryotes, three nuclear multisubunit RNA polymerases are essential for viability: RNA Polymerase I (Pol I), which synthesizes precursors for the three largest ribosomal RNAs, Pol II, which transcribes thousands of mRNAs and noncoding RNAs, and Pol III, which is required for 5S ribosomal RNA and tRNA biogenesis. Remarkably, plants have two additional nuclear RNA polymerases, Pol IV and Pol V, each composed of 12 subunits (1), at least seven of which are shared with Pol II, from which Pols IV and V evolved (2–5). In the siRNA-directed DNA methylation (RdDM) pathway, which primarily silences transposons, viruses and transgenes, Pols IV and V have non-redundant functions. Pol IV partners with RNA-DEPENDENT RNA POLYMERASE 2 (RDR2) to generate short double-stranded RNAs that serve as precursors for dicing into 24 nt siRNAs (6–8). These siRNAs are then incorporated into an Argonaute protein, primarily AGO4, and guide cytosine methylation and repressive chromatin modifications to loci transcribed by Pol V (9–11). Un-diced Pol IV and RDR2-dependent RNAs, and 21 nt siRNAs derived from degraded transposon mRNAs, are also implicated in guiding RNA-directed DNA methylation at sites of Pol V transcription (12–15).

Pols IV and V apparently have fewer constraints on their evolution than other polymerases, allowing their subunit compositions to vary (5) and their catalytic subunits to experience amino acid substitution rates that are ten to twenty times greater than for Pol II (3). More than 140 amino acid positions that are invariant in the catalytic subunits of Pols I, II, and III have diverged in Pols IV and V (Fig. S1A, S1B, Table S1) (16). These include substitutions and deletions within elements that are thought to be critically important for polymerase function, including the trigger loop and bridge helix (Fig. S1A) (17). In Pol II and other polymerases, conformational changes in the trigger loop and bridge helix result in the transition from an open state that allows nucleotide triphosphate (NTP) entry into the catalytic center to a closed state that has the NTP properly positioned relative to the 3' end of the nascent RNA chain, enabling phosphodiester bond formation (18, 19). Ratchet-like transitions between the open and closed conformations are thought to be linked to RNA translocation, affecting elongation rate as well as the accuracy (fidelity) of NTP incorporation (20, 21).

Despite the deletions and substitutions within the trigger loop, bridge helix, and other conserved domains, Pols IV and V have been shown to have RNA polymerase activity *in vitro* (22). However, the catalytic properties of Pol IV and Pol V, relative to Pol II, remain unknown. In this study, we conducted tests to compare several parameters of Pol II, Pol IV and Pol V catalytic activity. We find that Pols IV and V catalyze RNA synthesis more slowly than Pol II and we show that Pol IV transcription is error-prone. Surprisingly, Pol V is less error-prone than Pol II, at least in terms of misincorporating ribonucleoside triphosphates (rNTPs) mismatched to the DNA template. However, both Pol IV and Pol V exhibit reduced ability, relative to Pol II, to discriminate between ribonucleosides and deoxyribonucleosides. The implications of Pol IV and Pol V's enzymatic properties are discussed with respect to what is known about their functions.

## Results and Discussion

### Accuracy of Pol II, IV, and V transcription

We assessed polymerase fidelity (accuracy) using affinity-purified Pols II, IV or V to transcribe a 32 nt DNA template via extension of a 17 nt RNA whose 3’ half is complementary to the DNA template, yielding a 9 bp DNA-RNA hybrid (Fig. 1A). The RNA serves as a primer that can be elongated in a templated fashion by all three polymerases (22). The DNA templates were designed to have three identical nucleotides located immediately adjacent to the 3' end of the primer such that addition of a single, high-purity rNTP allows elongation of the RNA by 3 nt, to a length of 20 nt. Generation of 21 nt, or longer, RNAs indicates misincorporation of the NTP across from non-complementary nucleotides of the template. The assay depends on the use of high purity synthetic NTPs to avoid false-positive signals resulting from contamination by other NTPs, which can occur with NTPs purified from NTP mixtures (23).

**Figure 1.**
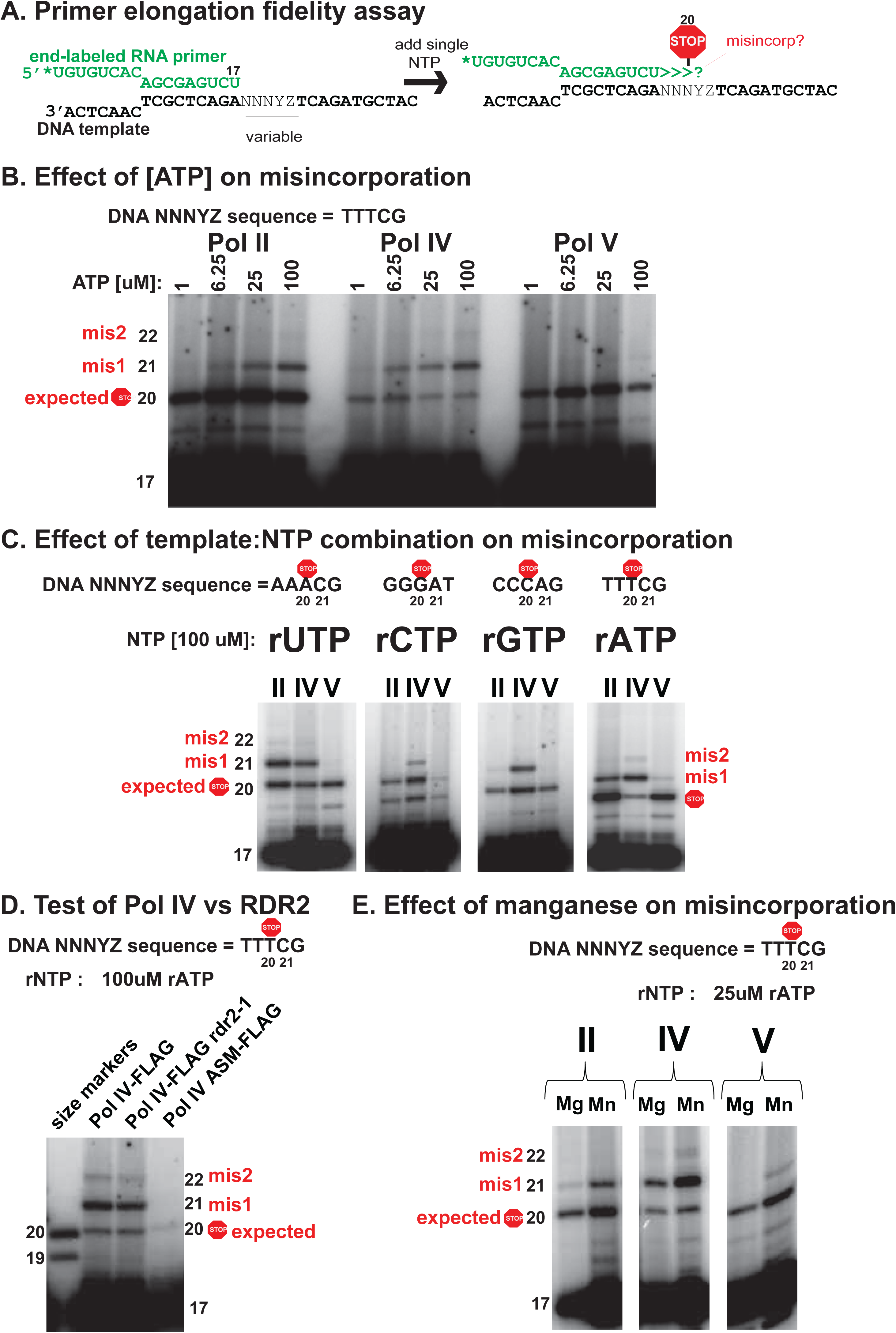
Pol IV and Pol V have altered fidelities relative to Pol II. *(A)* Design of the assay. *(B)* Pol II, IV, and V primer elongation products visualized by phosphorimaging following electrophoresis on a 15% PAGE gel. *(C)* Effect of NTP: DNA template combination on Pol II, IV, or V misincorporation. *(D)* Primer elongation fidelity assay comparing Pol IV immunoprecipitated from a line in which RDR2 co-immunoprecipitates with Pol IV (Pol IV-FLAG), an rdr2 null mutant line (Pol IV-FLAG, *rdr2-1*) or a line in which the NRPD1 transgene expresses an active site mutant (ASM-FLAG). *(E)* Effect of manganese in the transcription assay, in place of magnesium.

In primer elongation reactions involving adenosine incorporation, templated by Ts in the DNA, Pols II, IV, and V primarily synthesize the expected 20 nt RNA products when the ATP concentration is low (1 μM) (Fig. 1B). However, as the ATP concentration is increased (6.25 μM, 25 μM, or 100 μM), RNAs of 21 and 22 nt are synthesized as a result of misincorporating adenosine across from ensuing cytosine (misincorporation event 1; mis1) or guanosine (misincorporation event 2) bases of the DNA template. Comparing the ratio of properly arrested (20 nt) to misincorporation products (>20 nt), reveals that Pol IV generates the most misincorporation products, and Pol V the fewest (Fig. 1B). Subsequent tests comparing misincorporation frequency in the presence of 100 μM UTP, CTP, GTP or ATP confirmed that Pol IV is considerably more error-prone than Pols II or V, and that Pol V is the least error-prone of the three enzymes; this was true for all template and NTP combinations tested (Fig. 1C). The results also reveal that misincorporation varies considerably depending on the template-NTP combination, particularly for Pol II.

The RNA-dependent RNA polymerase, RDR2, physically associates with Pol IV (22, 24) and might plausibly contribute to NTP misincorporation. To test this possibility, we compared Pol IV isolated from wild-type *RDR2* plants, Pol IV isolated from a *rdr2-1* null mutant background, and Pol IV that is inactivated as a result of clustered point mutations in the Metal A site of the catalytic center, but expressed in a wild-type *RDR2* background (16). Pol IV isolated from the *rdr2-1* mutant misincorporated to the same extent as Pol IV isolated from wild type plants, whereas the Pol IV active site mutant lacked significant activity (Fig. 1D). Based on these controls, we conclude that Pol IV, and not RDR2, is responsible for the RNAs observed.

Magnesium ions are important for NTP positioning at the active site of RNA and DNA polymerases, such that substitution by bulkier manganese ions typically makes RNA and DNA polymerases error-prone (25, 26). We tested whether Pol IV and Pol V active sites are similarly sensitive to manganese. Indeed, substituting manganese for magnesium in the reaction buffer substantially increased nucleotide misincorporation by Pols IV and V, as for Pol II (Fig. 1E). These results suggest that the catalytic centers of Pols IV and V are similar to Pol II in their sensitivity to manganese ions.

To assess the contribution of nucleotide selectivity to Pol II, IV and V transcriptional fidelity, we compared the relative affinities of the enzymes for complementary versus non-complementary nucleotides during RNA elongation (18). For these assays, the 17 nt RNA primer was annealed to a template having CCCAG as the variable sequence downstream of the primer, such that addition of 0.1μM GTP resulted in polymerase-engaged elongation complexes that contain 20 nt RNAs (Fig 2A). Importantly, no misincorporation into 21 nt or longer products is detected using this low GTP concentration (compare to the 0 μM UTP reactions in Fig. 2B). The elongation complexes were then washed to remove templates not engaged by the resin-immobilized RNA polymerases, as well as free GTP, then incubated with UTP, the nucleotide complementary to the next template position, or ATP, which is non-complementary. Production of 21 nt or longer extension products was then monitored over a range of NTP concentrations, from 0 to 5 μM (5000 nM) for the complementary nucleotide (UTP) or 0 to 500 μM for the non-complementary nucleotide (ATP) (Fig. 2B). The band intensity for elongation products that are 21 nt or longer was then divided by the value of the total intensity for all products of 20 nt or longer and this ratio (expressed as %) was plotted versus UTP or ATP concentration (Fig. 2C, 2D). From these plots, we can obtain an estimated pseudo Km (Km*; considered a pseudo Km because we are not measuring velocity in these fixed-time reactions) as well as maximum incorporation (Imax) values, using linear regression to fit the data to the equation *y*=(*Imax*x*)/(*Km*+x*), as in the study of Wang *et al.* (18). Figure 2E shows the estimated Km*s for Pols II, IV, and V, for both complementary and non-complementary NTPs. All three polymerases have much higher (~1000-fold) estimated Km*s for the non-complementary (ATP) versus complementary nucleotide (UTP), indicating that the polymerases have much higher affinities for the correct versus incorrect NTP. Pol II and Pol V have similar Km*s for the complementary NTP (28 nM and 32 nM, respectively), but Pol IV has a much higher Km* (202 nM), suggesting decreased affinity for the correct NTP compared to Pols II or V (Fig. 2E). For the non-complementary nucleotide, Pol IV and Pol V have similar estimated Km*s (~112 uM) that are ~2 fold lower than for Pol II (252 μM) (Fig 2E).

**Figure 2.**
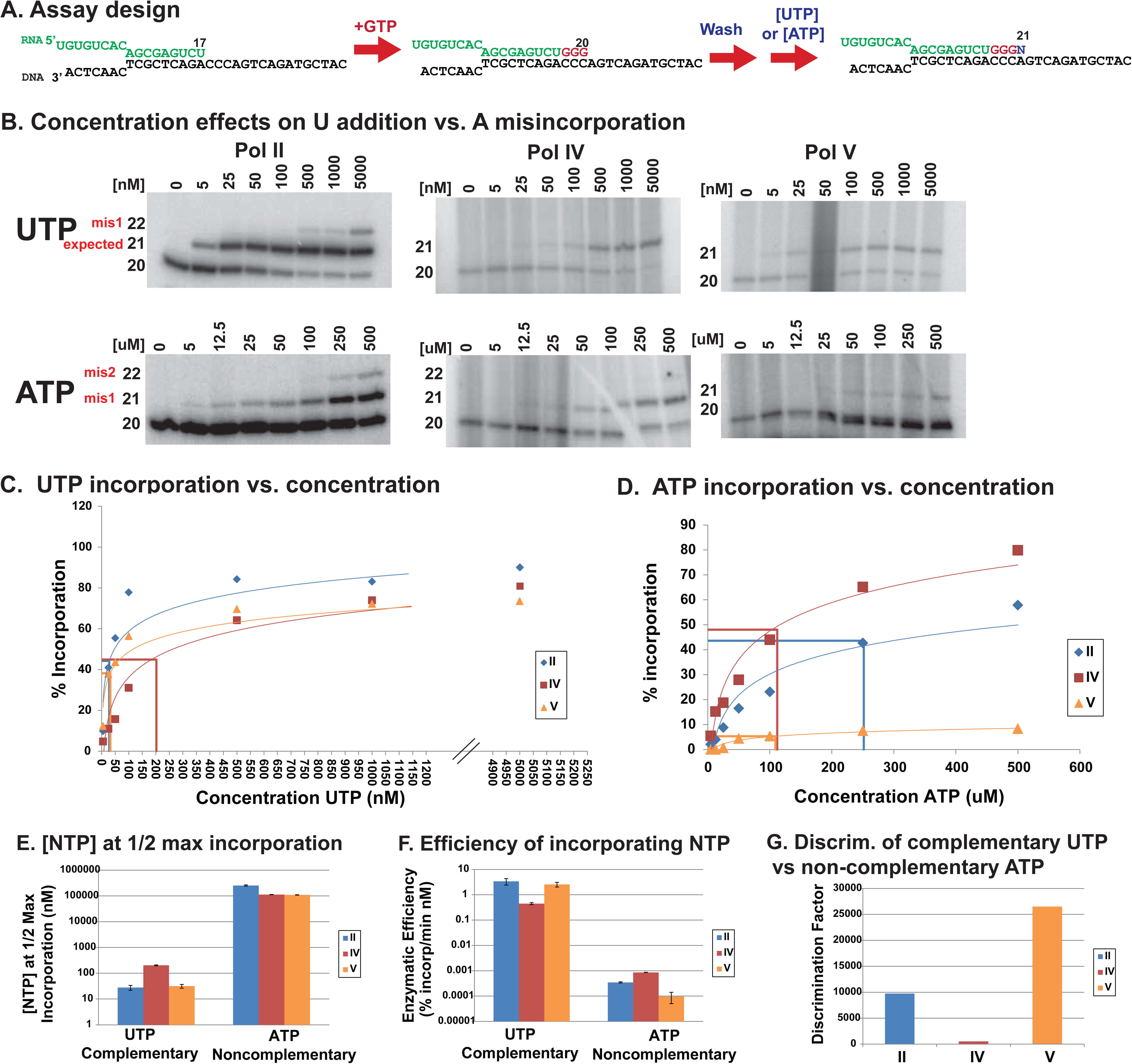
Differences in complementary versus non-complementary nucleotide incorporation contributes to differences Pol IV and Pol V fidelities, relative to Pol II. *(A)* Overview of the primer elongation assay. *(B)* Representative gels showing the incorporation of the complementary nucleotide (UTP) by Pols II, IV, and V over a range of UTP concentrations. *(C)* Percent incorporation, calculated as the intensity of products of 21 and 22nts divided by the total intensity of products 20nts or longer) is plotted versus nucleotide concentration, with linear regression used to fit the curves to the equation *y=(Imax*x)/(Km*+x). (D)* Percent incorporation (intensity of 21 and 22 nt products divided by the total intensity of products 20 nts and longer) plotted versus nucleotide concentration, with linear regression used to fit the curves to the equation *y=(Imax*x)/(Km*+x). (E)* Km* values (nucleotide concentration at ½maximum incorporation (Imax)) of Pols II, IV, and V for complementary and non-complementary NTPs. Data is representative of two replicate experiments (see Fig. S2). *(F)* Efficiency of nucleotide incorporation (*Imax/Km\**). *(G)* Discrimination factor, calculated as complementary NTP efficiency/non-complementary NTP efficiency.

Pol IV’s decreased affinity for the correct NTP and increased affinity for a non-complementary NTP is consistent with Pol IV's propensity for misincorporation, as shown in Figure 1. Pol V, on the other hand, has a Km* for the complementary nucleotide that is similar to Pol II, but has a lower Km* for the non-complementary nucleotide, suggesting that Pol V has a higher affinity for the wrong nucleotide compared to Pol II. This was unexpected given that Pol V produces fewer misincorporation products than Pol II in the fidelity assays of Figure 1. The explanation for this apparent paradox comes from considering enzymatic efficiency, which in conventional Michaelis-Menten analyses is estimated by dividing Vmax (the concentration of substrate at which the enzymes active site is saturated) by the Km, substituted in our case by Imax and Km* (Fig. 2F). Pol II and Pol V are similarly efficient at incorporating the complementary nucleotide, whereas Pol IV is relatively inefficient. By contrast, Pol IV is the most efficient at incorporating the non-complementary nucleotide, whereas Pol V is least efficient. Collectively, these experiments indicate that although Pol V has a higher affinity for the noncomplementary NTP compared to Pol II (Fig. 2E), it has a lower Imax (Fig. 2D) such that the overall efficiency of incorporating a wrong nucleotide is lower than for Pol II (Fig 2F). Pol V's 3-fold increased propensity to incorporate correct versus incorrect nucleotides, compared to Pol II, and Pol IV’s 19-fold decreased ability compared to Pol II (Fig 2G), are consistent with the misincorporation results of Figure 1.

We tested whether nucleotide misincorporation is enhanced by cytosine methylation, as this has been postulated to occur and possibly contribute to Pol IV termination (7). Comparing methylated and unmethylated DNA template oligonucleotides in primer elongation experiments (Fig. 3A), we observed no increase in misincorporation of ATP, UTP or CTP caused by methylation of template cytosines, in either 5' ^me^CHH or 5' ^me^CG sequence contexts, compared to unmethylated template cytosines (Fig. 3B, 3C). These results suggest that misincorporation or rNTPs at methylcytosine positions of the template is not an intrinsic property of Pols II, IV or V.

**Figure 3.**
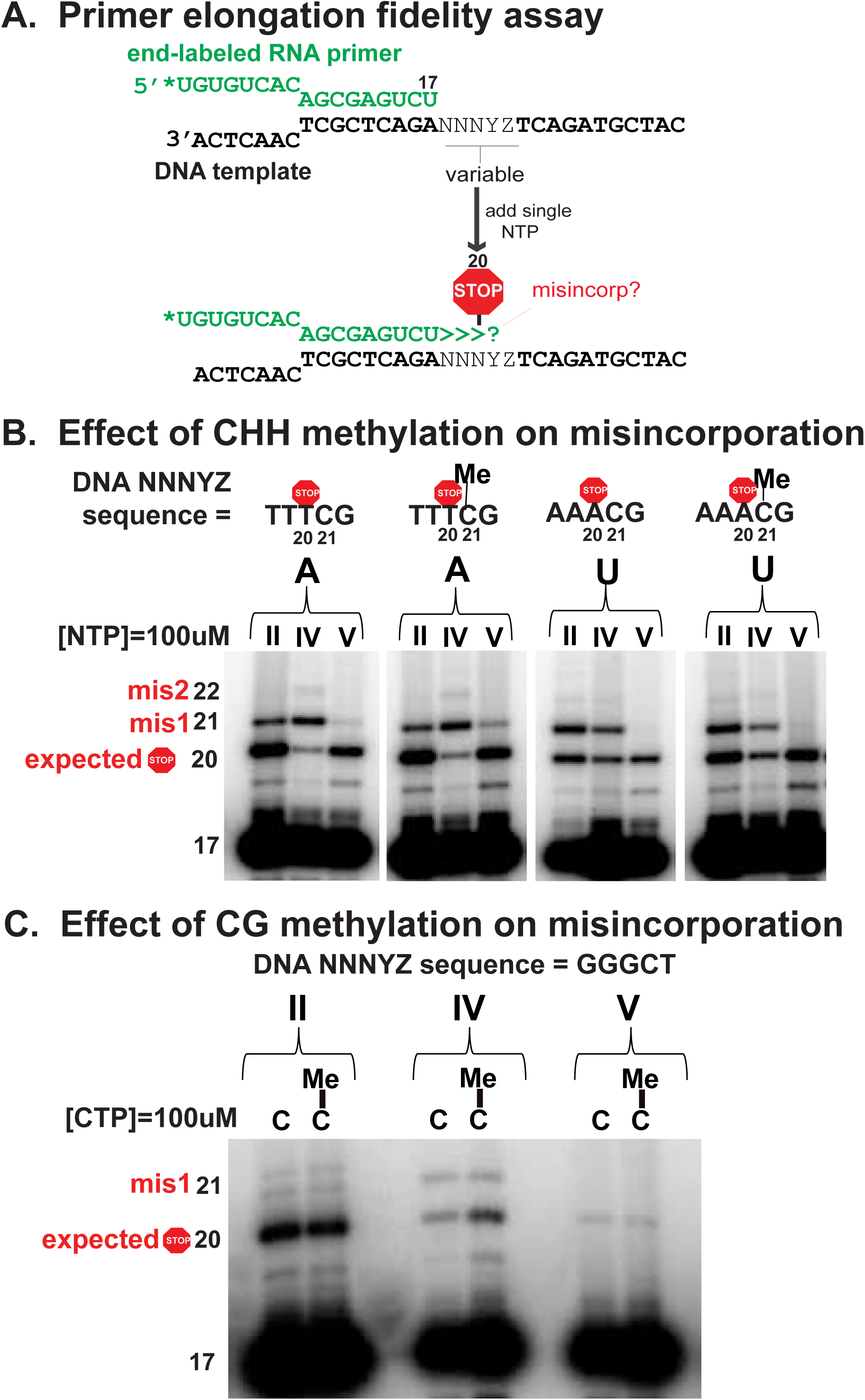
Effect of DNA template cytosine methylation on Pol II, IV, or V transcriptional fidelity. *(A)* Overview of primer elongation assay. *(B)* DNA template CHH methylation tests, for ATP or UTP misincorporation *(C)* DNA template CG methylation tests, for CTP misincorporation.

### Discrimination between rNTPs and dNTPs

To test the abilities of Pol IV and Pol V to discriminate between ribo and deoxyribo-nucleoside triphosphates, primer elongation experiments were conducted in the presence of 100% rNTP, 100% dNTP or a 50:50% mix of rNTP and dNTP (Fig. 4A). Primers that are elongated by incorporating dNTPs migrate faster than rNTP-elongated primers when subjected to electrophoresis on 15% denaturing polyacrylamide sequencing gels (Fig. 4B). When provided a 50:50 mix of rNTP and dNTP, Pol II makes products whose mobility is the same as when only rNTP is provided, demonstrating a strong preference for rNTPs over dNTPs (Fig. 4B). Pol IV prefers rNTPs, but dNTP elongation products are also detected in reactions containing equal amounts of both types of NTP(Fig. 4B). Pol V, surprisingly, displays similar incorporation of rNTPs or dNTPs when either is provided alone, and preferentially incorporates the dNTPs when provided with a 50:50 mix (Fig. 4B). These results show that Pols IV and V have a reduced ability, compared to Pol II, to discriminate between rNTPs and dNTPs. In order to better understand the basis for this loss of discrimination, we determined the affinities of Pols II, IV, and V for a dNTP in the same way rNTP affinity was assessed in Figure 2 (Fig. S3A). We found that Pols IV and V have increased affinities, relative to Pol II, for dNTP (Fig S3B-G).

**Figure 4.**
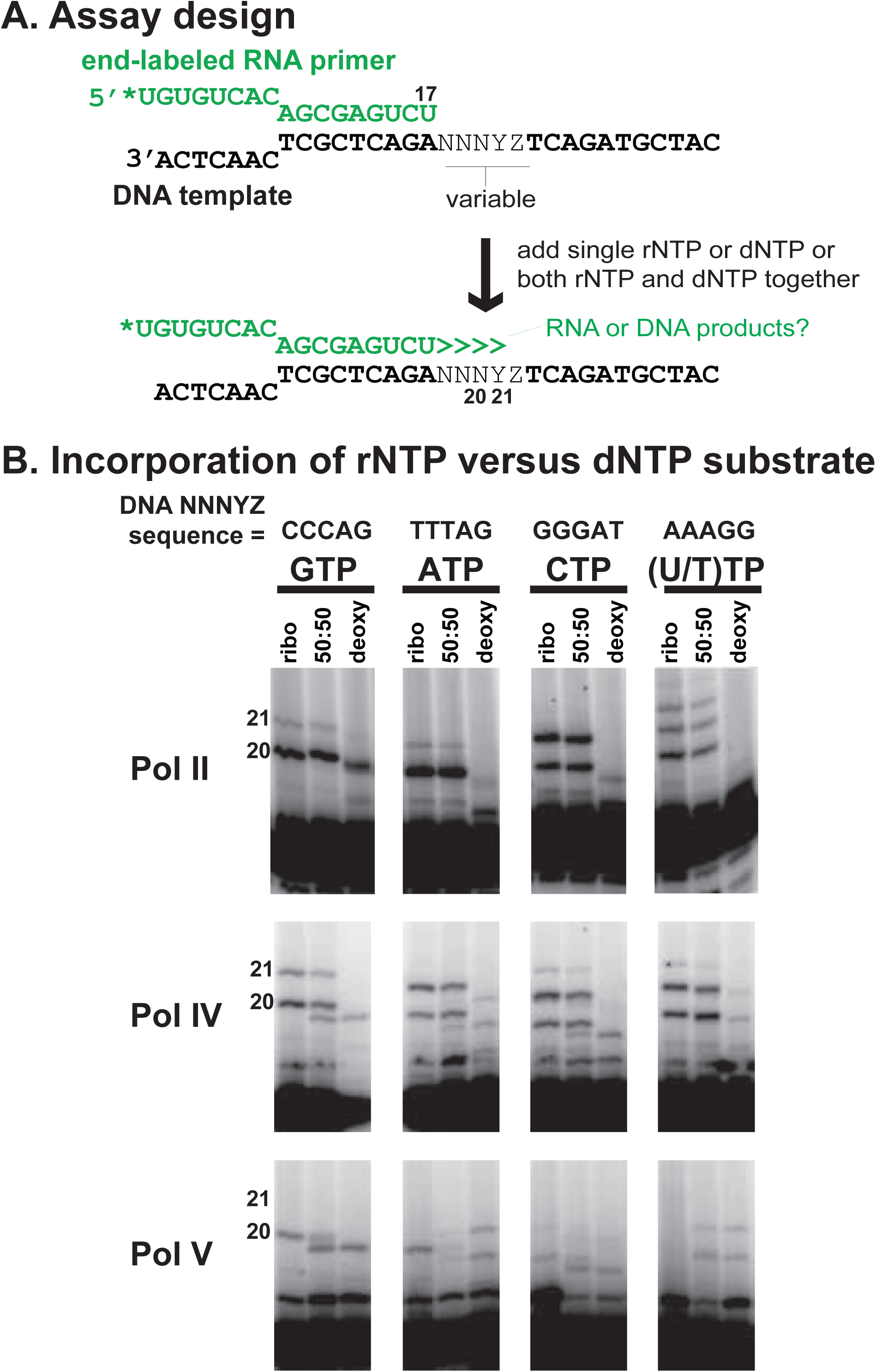
Pols IV and V misincorporate dNTPs to a greater extent than Pol II. *(A)* Assay design. *(B)* Primer elongation assays in the presence of only rNTP, only dNTP, or a 50:50 rNTP:dNTP mix.

### Rates of nucleotide incorporation

Relative rates of nucleotide incorporation for Pols II, IV and V were assessed by elongating a 17 nt primer to 20 nt RNA using a low concentration of GTP as in Fig. 2 (Fig. 5A), then conducting a time-course of 21 nt RNA production upon addition of 500 nM UTP (Fig. 5B). The percentage of 21 nt product, relative to total 20 + 21 nt product, was plotted against time (Fig. 5C) and the data were fitted to the equation *c(t)* = *A x (1 – exp[-k x t])* to calculate the rate constant, k, as described in Sydow *et al.* (23). Pol IV and Pol V both have decreased rates of nucleotide incorporation relative to Pol II (Fig. 5D). Pol IV is the slowest of the three enzymes, with a rate that is ~6 times slower than Pol II (k values of 0.06 versus 0.38, respectively). Pol V is 3 times slower than Pol II (k values of 0.13 versus 0.38, respectively), yet still about twice as fast as Pol IV (Fig 5D).

**Figure 5.**
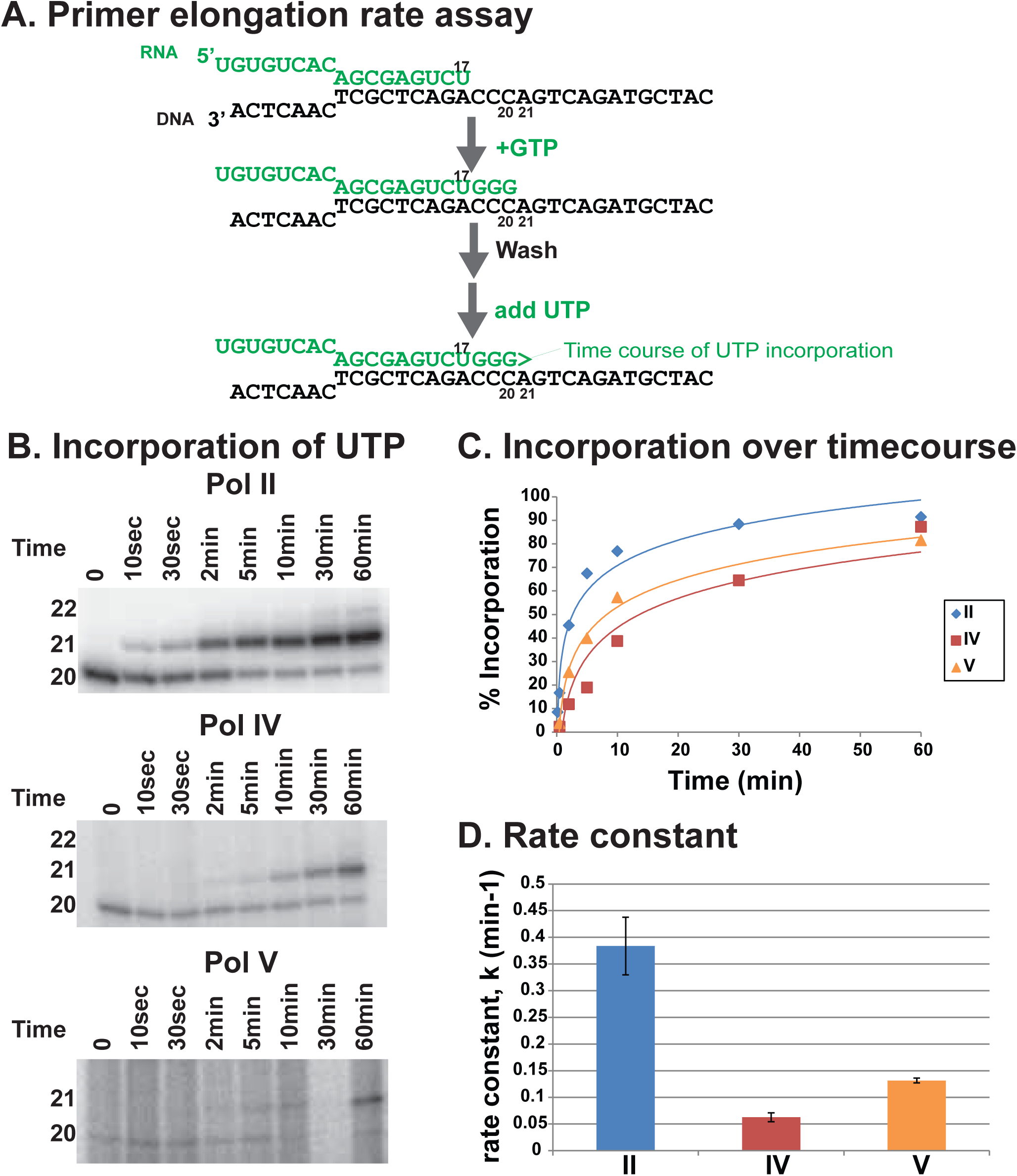
Relative nucleotide incorporation rates of Pols II, IV and V. *(A)* Assay design. *(B)* Representative gels showing the incorporation of UTP by Pols II, IV, and V versus time (UTP concentration = 500 nM). *(C)* Percent incorporation (intensity of elongation products divided by total products) versus time, with linear regression used to fit the curves to the equation *c(t) = A x (1 – exp[-k x t]). (D)* Elongation rate constants for Pols II, IV and V. Data is representative of two replicate experiments (see Fig. S4).

### Transcription error rates in *de novo* synthesized RNAs

To test whether differences in Pol II and Pol IV fidelity observed using defined template oligonucleotides, and primer extension with only one or two NTPs, are also observed for transcripts initiated *de novo* in a primer-independent manner, in the presence of all four NTPs, we sequenced RNAs generated by Pol II and Pol IV using single-stranded, circular bacteriophage M13mp18 DNA as the template. Pols II and IV initiate at more than two thousand distinct start sites within the ~7.2 kb M13 genome sequence, providing a diverse set of transcripts (6). The resulting Pol II and Pol IV transcripts were subjected to “circle sequencing” (Fig 6A) (27). In this method, the 5' and 3' ends of transcripts are ligated to form circles prior to cDNA synthesis using SuperScript III Reverse Transcriptase (Thermo Fisher Scientific). This enzyme has strand displacement activity, allowing it to reverse-transcribe a RNA circle multiple times, producing a cDNA concatamer consisting of multiple DNA copies of the original RNA template. True transcription errors present in the RNA will be present in each repeat of the concatamer whereas sporadic errors introduced by the reverse transcriptase, PCR polymerase, or sequencing polymerase will not, allowing these sources of error to be discriminated from one another. Variations of this method have been used to identify genetic variants in RNA viral populations (28, 29), as well as to measure the transcriptional fidelities of bacterial RNA polymerases (30).

**Figure 6.**
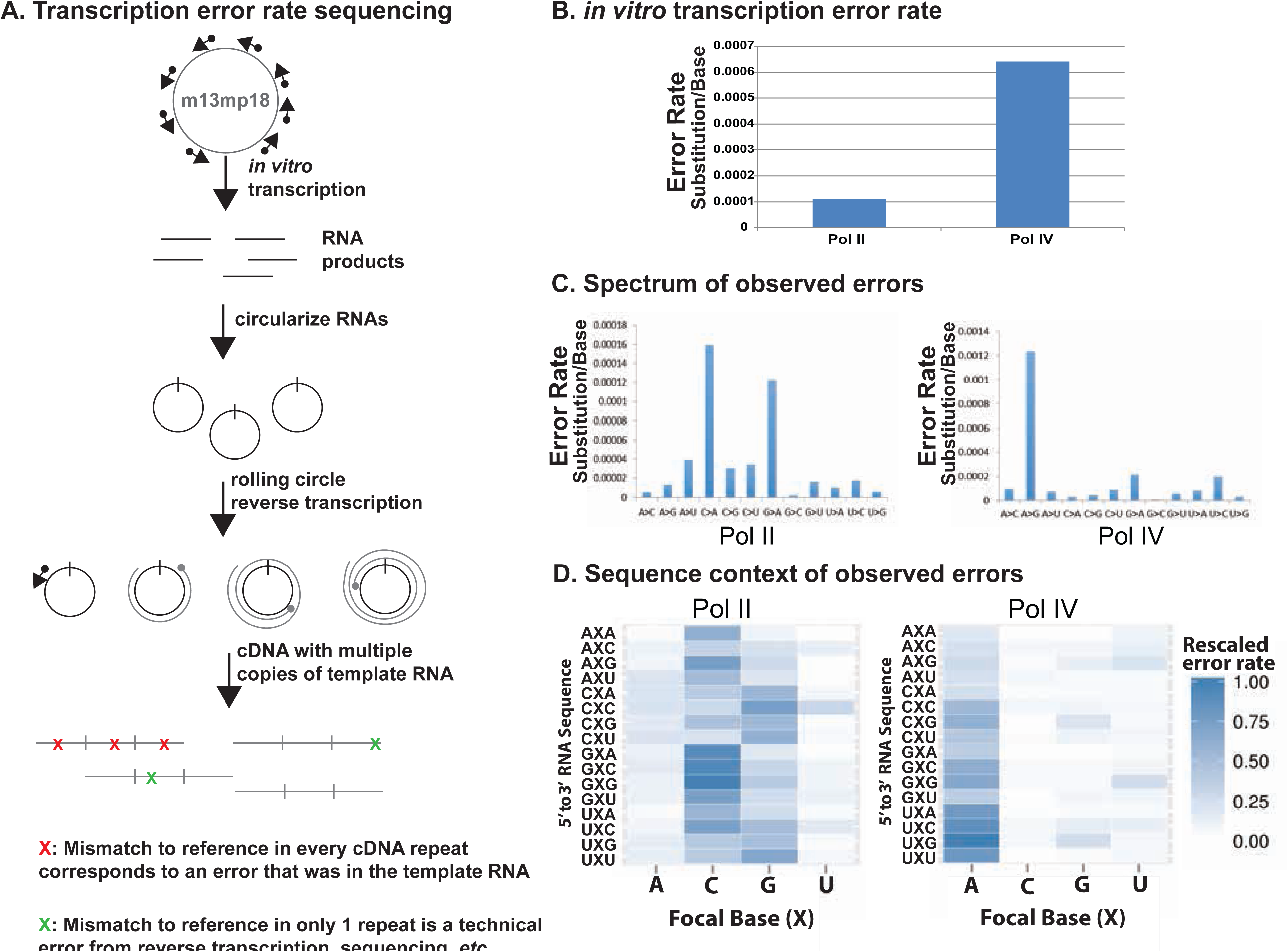
Pol IV exhibits increased error rate and an altered pattern of misincorpration relative to Pol II in RNAs initiated *de novo* (primer-independent), using single-stranded M13 DNA as the template. *(A)* Assay design for circle sequencing of cDNA concatamers. *(B)* Overall error rates of Pol II and Pol IV, expressed as substitution frequency per base sequenced. *(C)* Error rates across the spectrum of misincorporation events generated by Pol II or Pol IV, expressed as substitution frequency per base sequenced. *(D)* 5’ and 3’ sequence contexts for RNA nucleotide positions at which errors were observed. Focal Base denotes the nucleotide where misincorporation occurred.

Using the circular sequencing approach, transcription error rates for RNAs generated by Pol II or Pol IV using the M13 template were calculated as the number of errors divided by the total number of nucleotides sequenced. Pol IV's *in vitro* error rate was found to be roughly six times greater than the transcription error rate for Pol II (6.4 × 10^−4^ and 1.1 × 10^−4^, respectively) (Fig 6B). These results support our findings using primer elongation with single nucleotides, which also showed that Pol IV has an increased propensity for misincorporation relative to Pol II.

The observed Pol II *in vitro* error rate is consistent with a previously estimated *in vitro* error rate for Pol II isolated from wheat germ, which ranged from 10^−4^ to 10^−6^ depending on the combination of NTP and DNA template (31). The Pol II error rate in our study varied with NTP/template combination (Fig. 6C) and the sequence context of flanking nucleotides (Fig 6D). As in the study by de Mercoyrol *et al*, Pol II misincorporation of A opposite dC or dG in the template occurs at a frequency that is an order of magnitude higher than for other RNA:template combinations (Fig 6C).

Interestingly, the misincorporation spectrum differs between Pol II and Pol IV, with Pol IV exhibiting a strong bias for misincorporating G opposite dT in the template (Fig 6B). Pol II and IV also differ in the sequence context of misincorporation events. Figure 6D depicts the nucleotides flanking misincorporated nucleotides at the focal base (central) position of nucleotide triplets, with darker color indicating a higher degree of misincorporation. Pol II displays a preference for misincorporating after C or G; whereas, Pol IV exhibits a trend for misincorporating after U (Fig 6D).

## Discussion

Collectively, our investigation of Pol IV and V catalytic properties provides initial insights into the functional characteristics of the enzymes. Pol IV and Pol V differ from Pol II, and from one another, in a number of enzymatic properties, including accuracy and catalytic rate. Pol IV is the slowest of the three enzymes, and is also the most error-prone. Pol IV/RDR2-dependent precursor RNAs are only ~30-40 nt in length, just long enough to encode 24 nt siRNAs (6, 7), such that Pol IV may not need to be fast. And because 24 nt siRNAs primarily guide the silencing of transposons whose family members are not necessarily identical, being error-prone may be tolerated, and even beneficial. The accuracy of Pol IV transcription would, however, affect whether an siRNA binds a target RNA (or DNA) strand with perfect or imperfect complementarity, potentially influencing whether AGO4 might slice, or just bind, Pol V target transcripts (32). Evidence suggests that slicing activity is important at some, but not all, RdDM loci (33).

Pol V makes longer transcripts than Pol IV. Although their precise size remains undefined, RT-PCR analyses indicate that they can be 200 nt or more (34–36). Our results indicate that Pol V transcription is also highly accurate, suggesting that accuracy is important for Pol V transcript function. Pol V makes RNAs at loci to be silenced by RNA-directed DNA methylation, and its transcripts are thought to provide scaffolds for the binding of siRNA-AGO complexes that then recruit additional chromatin modifying activities (11, 34), consistent with studies in fission yeast and other organisms (37). A need for precise basepairing between siRNAs and Pol V transcripts might be a selective pressure for maintaining Pol V fidelity. A recent study has suggested that siRNA-AGO4 complexes may also bind directly to DNA at Pol V-transcribed loci (38). If the act of transcription is all that is needed for Pol V to function, it is not clear why Pol V would need to generate RNAs that faithfully match the sequence of transcribed loci. One possibility might be that siRNAs first bind Pol V transcripts prior to binding the corresponding DNA sequence. Another possibility is that Pol V transcripts are used to generate R-loops at transcribed loci, thereby enabling siRNA-AGO interactions with the displaced strand (39), with precise basepairing of the RNA and DNA being potentially important.

Some substitutions of ultra-conserved amino acids in Pols IV and V occur at positions known to affect RNA polymerase fidelity. For example, mutation of N479 in the *S. cerevisiae* RNA Pol II largest subunit, Rpb1 results in reduced discrimination between NTPs and dNTPs (18). Both Pol IV and Pol V have substitutions at the homologous position, consistent with their incorporation of dNTPs to a greater extent than Pol II. However, the details of altered rNTP:dNTP discrimination differs in the yeast N479 Pol II mutant versus Pols IV and V. The N479S Pol II mutant has decreased affinity for rNTPs, rather than increased affinity for the dNTPs (18). In contrast, Pol IV and Pol V have increased dNTP affinity (Figure S3). Additional diverged amino acids of Pols IV and V presumably contribute to these differences.

It is not clear whether incorporating dNTPs into Pol IV or Pol V transcription products is biologically meaningful. Outside of S phase of the cell cycle, rNTP concentrations are expected to far exceed dNTP concentrations. Misincorporation of dNTPs into RNA by a T7 RNA polymerase mutant has been found to block translation of the dNTP-carrying transcripts (40). Therefore, one could speculate that incorporating dNTPs into Pol IV or Pol V transcripts could potentially help ensure that the transcripts are not translated, but this seems unnecessary for Pol IV and Pol V RNAs acting in the nucleus. Incorporation of dNTPs might potentially affect binding or processing of Pol IV or Pol V transcripts. Similar to misincorporation of a non-complementary base, misincorporation of a dNTP reduces the likelihood of adding a subsequent nucleotide (18). Therefore, incorporation of dNTPs into Pol IV and V transcripts might also contribute to stalling or termination. Pol IV and Pol V have also been implicated in DNA double-strand break repair, suggesting that an ability to synthesize products incorporating dNTPs could be important for this process (41).

The trigger loop within the largest subunit of yeast RNA Pol II is thought to play an important role in transcriptional fidelity. Amino acids within the trigger loop (Leu1081, Gln1078, His1083, and Asn 1082) contact the base, phosphate, and ribose of an incoming NTP to facilitate precise positioning and catalysis (18). Pols IV and V have amino acid substitutions at three of these four positions (Fig. S1A). In addition, *A. thaliana* Pol IV, which has reduced accuracy relative to Pol II, has diverged at a trigger loop position whose mutation in the yeast Pol II largest subunit, Rpb1 (position E1103) results in increased NTP misincorporation (20). The E1103G mutation is thought to destabilize the active site open conformation, causing increased misincorporation due to greater sequestration of non-complementary nucleotides within the closed conformation (20). It has been suggested that other conditions that reduce Pol II fidelity, such as deletion of the ninth subunit, or the presence of manganese, similarly promote a closed trigger loop conformation (18, 20, 42, 43). Opening and closing of the trigger loop is also important for elongation, such that mutations alter the catalytic rate (44, 45). Given their altered fidelities (Figs. 1,2, and 4), rates of nucleotide incorporation (Fig. 5), and extensive sequence divergence (or deletion) in the trigger loop region (Fig. S1), we speculate that the Pol IV active center may naturally adopt a more closed structure, whereas the Pol V active center may resemble the Pol II open conformation. Such speculations underscore a need for additional mechanistic studies of Pol IV and Pol V transcription, which would be benefitted significantly by high-resolution structural models for the enzymes.

## Materials and Methods

### Protein alignments

Amino acid sequences for the largest and second-largest subunits of RNA polymerases I through V, from multiple species, were obtained from the National Center for Biotechnolgy Information (https://www.ncbi.nlm.nih.gov/). Sequences were aligned using Clustal Omega (46, 47).

### RNA Polymerase affinity purification

Pols II, IV, and V were immunoprecipitated from leaf tissue of 3 week old transgenic lines expressing FLAG epitope-tagged RNA polymerase subunits, NRPB2-FLAG (Pol II), NRPD1-FLAG (Pol IV), or NRPE1-FLAG (Pol V), as previously described (22). Leaf tissue was ground to a powder in liquid nitrogen using a mortar and pestle and then resuspended in 3.5 mL extraction buffer (20 mM Tris-HCl, pH 7.6, 300 mM sodium sulfate, 5 mM magnesium sulfate, 5 mM DTT, 1 mM PMSF, 1% plant protease inhibitor (Sigma) per gram of tissue. The homogenate was subjected to centrifugation at 16,000 x g for 15 minutes, 4°C. The supernatant was collected and subjected to a second round of centrifugation using the same conditions. 50 μL anti-FLAG agarose resin (Sigma) was added to each lysate and incubated at 4°C for 3 hours on a rotating mixer. Resin was washed twice with 10 mL wash buffer (20 mM Tris-HCl, pH 7.6, 300 mM sodium sulfate, 5 mM magnesium sulfate, 5 mM DTT, 1 mM PMSF, 0.5% IGEPAL CA-630 detergent) and once with 10 mL CB100 (25 mM HEPES-KOH, pH 7.9, 20% glycerol, 100 mM KCL, 1 mM DTT, 1 mM PMSF). Immunoprecipitated polymerases were used immediately in *in vitro* transcription reactions. Pol II immunoprecipitated from 1 gram of leaf tissue expressing NRPB2-FLAG was sufficient for 20 *in vitro* transcription reactions, whereas 4 grams of NRPD1-FLAG or NRPE1-FLAG leaf tissue was needed for single reactions.

### *in vitro* transcription using oligonucleotide templates

*in vitro* transcription reactions were conducted using a 17 nt RNA primer hybridized to various 32 nt ssDNA oligos, as previously described (22), with minor modifications. RNA primers (2 μM) were end-labelled using T4 polynucleotide kinase and Y-^32^P-ATP, and excess Y-^32^P-ATP was removed using Performa spin columns according to the manufacturer’s protocol (Edge Bio). RNA-DNA hybrid templates were generated by combining equimolar amounts of end-labelled RNA primer and unlabeled DNA template in 1X annealing buffer (100 mM potassium acetate, 30 mM HEPES-KOH, pH 7.5), placed in a boiling water bath, and allowed to cool to room temperature. Template sequences are provided in Table S2.

50 μL of resuspended, washed polymerase, still bound to the immunoprecipitation resin, was used for each transcription reaction. Washed NRPB2-FLAG resin (Pol II) was resuspended in 1 mL CB100 buffer, enough for 20 reactions. Washed Pol IV and Pol V resins were resuspended in a final volume of 50 μL per transcription reaction. 50 μL of 2X transcription reaction mix containing 21.7 μl of the 250 nM, annealed RNA-DNA template solution, 100 μΜ (unless otherwise noted) high purity rNTPs (GE Healthcare), 120 mM ammonium sulfate, 40 mM HEPES-KOH pH 7.9, 24 mM magnesium sulfate, 20 uM zinc sulfate, 20 mM DTT, 20% glycerol, and 1.6 U/μL Ribolock (Thermo Fisher Scientific) was then added. In rNTP:dNTP discrimination experiments, the total NTP concentration was 100 μΜ. The high purity NTPs (GE Healthcare) used are free of other NTPs or RNAse activity. NTPs that are synthesized, not purified from a mixture of nucleotides, are not susceptible to cross-contamination during manufacture (23), an important consideration for misincorporation assays. Reactions were incubated at room temperature for 1 hour on a rotating mixer. Reactions were desalted using Performa spin columns according to the manufacturer’s protocol (Edge Bio), then precipitated with 1/10 volume 3M sodium acetate, 20 μg GlycoBlue (Thermo Fisher Scientific) and an equal volume of isopropanol. Pellets were resuspended in 5 μL RNA gel loading dye (47.5% formamide, 0.01% SDS, 0.01% bromophenol blue, 0.005% xylene cyanol, 0.5 mM EDTA), heated at 70°C for 3 minutes, and subjected to electrophoresis on a 15% polyacrylamide, 7M urea sequencing gel.

### Km* and rate assays

Km* and rate assays were performed similar to *in vitro* transcription assays described above, with minor modifications. Transcription was initiated by adding 50 μL 2X transcription reaction mix containing 0.1 μΜ of the first nucleotide complementary to the template (rGTP), and incubation for 20 min. The elongation complexes were then washed with 800 μL 1X transcription buffer lacking NTPs (60 mM ammonium sulfate, 20 mM HEPES-KOH pH 7.9, 12 mM magnesium sulfate, 10 μΜ zinc sulfate, 10 mM DTT, 10% glycerol). 0.8 U/μL of RNase inhibitor was added to washed elongation complex resin. For Km* reactions, 100 μL washed resin was distributed to 1.5 mL tubes containing appropriate volumes of the next complementary nucleotide, or a non-complementary nucleotide, to achieve the substrate concentration being tested for that nucleotide. Reactions were incubated at room temperature, on a rotating mixer, for 30 minutes. For rate assays, 500 nM of the complementary nucleotide (UTP) was added to the washed elongation complexes and the reactions were incubated at room temperature for the range of times indicated in the figure. Reactions were cleaned and analyzed as described for *in vitro* transcription assays above.

### Error rate determination for *de novo*-initiated M13 template transcripts

To determine error rates for Pol II or Pol IV transcripts generated from M13mp18 single-stranded DNA (Bayou Biolabs) as the template, transcription reactions were conducted as previously described, but in a transcription reaction mix consisting of 7.5 nM M13mp18 (Bayou Biolabs), 1 mM ATP, 1 mM GTP, 1 mM CTP, 1 mM UTP, 60 mM ammonium sulfate, 20 mM HEPES-KOH pH 7.9, 12 mM magnesium sulfate, 10 uM zinc sulfate, 10 mM DTT, 10% glycerol, and 0.8 U/μL Ribolock (Thermo Fisher Scientific). Reaction products were purified using Performa spin columns according to the manufacturer’s protocol (Edge Bio), then precipitated with 1/10 volume 3M sodium acetate, 20 μg GlycoBlue (Thermo Fisher Scientific) and an equal volume of isopropanol. Pellets were resuspended in 5 μL nuclease-free water and DNase treated using a Turbo DNA-free kit (Thermo Fisher Scientific) according to the manufacturer’s protocol. RNAs were then treated with RNA 5’ pyrophosphohydrolase (RppH from New England Biolabs) to convert 5’-end triphosphates to 5’ monophosphates. Reactions were cleaned with Oligo Clean & Concentrator columns (Zymo Research) according to the manufacturer’s protocol. Without any fragmentation, RNAs were circularized with RNA ligase 1 (NEB, M0204S) according to the manufacture’s guidelines. Circularized RNA templates were then reverse transcribed in a rolling-circle reaction according to the protocol described by Acevedo et al, with the exception that the incubation time at 42 °C was extended from 2 minutes to 20 minutes (28, 29). Second strand synthesis and the remaining steps for the library preparation were then performed using a NEBNext Ultra RNA Library Pre Kit for Illumina (E7530L) and the NEBNext Multiplex Oligos for Illumina (E7335S, E7500S), according to the manufacturer’s protocols. A size selection for amplified products longer than 300nt was performed before sequencing and 300nt single-end reads were then generated using an Illumina Hiseq instrument. Following the autocorrelation-based method and Bayesian approach described by Lou and Hussman *et al*, the structure of repeats within a read was identified and the consensus sequence of a repeat was constructed (27). Because of the random-priming approach used for rolling-circle reverse transcription, the 5’ end of the consensus sequence can be any nucleotide of the circularized RNA template. To reorganize the consensus sequence and make the ends correspond to the 5’ and 3’ of the original RNA transcript, we first constructed a tandem duplicate of the consensus sequence and mapped it back to the M13mp18 reference by BWA (48). Therefore, the longest continuous mapping region of the duplicated consensus sequence corresponds to the original RNA transcript. Since the mapping results can be ambiguous at the first and last few nucleotides, we excluded the 4 nucleotides at each end of the reorganized consensus sequence prior to subsequent transcript analyses to minimize potential false positives. The reconstructed consensus sequence was then mapped to the M13mp18 reference sequence, with transcription errors called for mismatches present in tandem copies of the RNA, and a frequency of mismatch no larger than 1%.

## Acknowledgements

This work was supported by funds to CSP as an Investigator of the Howard Hughes Medical Institute and Gordon and Betty Moore Foundation and from grants GM077590 (CSP) and GM036827 (ML) from the National Institutes of Health. MM received support from National Institutes of Health training grant, T32GM007757.

## References

1. Ream TS, et al. (2009) Subunit compositions of the RNA-silencing enzymes Pol IV and Pol V reveal their origins as specialized forms of RNA polymerase II. Mol Cell 33(2): 192–203.

2. Huang Y, et al. (2015) Ancient Origin and Recent Innovations of RNA Polymerase IV and V. Mol Biol Evol 32(7): 1788–1799.

3. Luo J & Hall BD (2007) A multistep process gave rise to RNA polymerase IV of land plants. J Mol Evol 64(1):101–112.

4. Tucker SL, Reece J, Ream TS, & Pikaard CS (2010) Evolutionary history of plant multisubunit RNA polymerases IV and V: subunit origins via genome-wide and segmental gene duplications, retrotransposition, and lineage-specific subfunctionalization. Cold Spring Harb Symp Quant Biol 75:285–297.

5. Haag JR, et al. (2014) Functional Diversification of Maize RNA Polymerase IV and V Subtypes via Alternative Catalytic Subunits. Cell Rep 9(1):378–390.

6. Blevins T, et al. (2015) Identification of Pol IV and RDR2-dependent precursors of 24 nt siRNAs guiding de novo DNA methylation in Arabidopsis. Elife 4:e09591.

7. Zhai J, et al. (2015) A One Precursor One siRNA Model for Pol IV-Dependent siRNA Biogenesis. Cell 163(2):445–455.

8. Li S, et al. (2015) Detection of Pol IV/RDR2-dependent transcripts at the genomic scale in Arabidopsis reveals features and regulation of siRNA biogenesis. Genome Res 25(2):235–245.

9. Matzke MA & Mosher RA (2014) RNA-directed DNA methylation: an epigenetic pathway of increasing complexity. Nat Rev Genet 15(6):394–408.

10. Wendte JM & Pikaard CS (2016) The RNAs of RNA-directed DNA methylation. Biochim Biophys Acta.

11. Wierzbicki AT, Ream TS, Haag JR, & Pikaard CS (2009) RNA polymerase V transcription guides ARGONAUTE4 to chromatin. Nature genetics 41(5):630–634.

12. Yang DL, et al. (2016) Dicer-independent RNA-directed DNA methylation in Arabidopsis. Cell Res 26(1):66–82.

13. Ye R, et al. (2016) A Dicer-Independent Route for Biogenesis of siRNAs that Direct DNA Methylation in Arabidopsis. Mol Cell 61(2):222–235.

14. Nuthikattu S, et al. (2013) The initiation of epigenetic silencing of active transposable elements is triggered by RDR6 and 21-22 nucleotide small interfering RNAs. Plant Physiol 162(1):116–131.

15. McCue AD, et al. (2015) ARGONAUTE 6 bridges transposable element mRNA-derived siRNAs to the establishment of DNA methylation. EMBO J 34(1):20–35.

16. Haag JR, Pontes O, & Pikaard CS (2009) Metal A and metal B sites of nuclear RNA polymerases Pol IV and Pol V are required for siRNA-dependent DNA methylation and gene silencing. PLoS One 4(1):e4110.

17. Landick R (2009) Functional divergence in the growing family of RNA polymerases. Structure 17(3):323–325.

18. Wang D, Bushnell DA, Westover KD, Kaplan CD, & Kornberg RD (2006) Structural basis of transcription: role of the trigger loop in substrate specificity and catalysis. Cell 127(5):941–954.

19. Vassylyev DG, et al. (2007) Structural basis for substrate loading in bacterial RNA polymerase. Nature 448(7150): 163–168.

20. Kireeva ML, et al. (2008) Transient reversal of RNA polymerase II active site closing controls fidelity of transcription elongation. Molecular Cell 30(5):557–566.

21. Feig M & Burton ZF (2010) RNA polymerase II with open and closed trigger loops: active site dynamics and nucleic acid translocation. Biophys J 99(8):2577–2586.

22. Haag JR, et al. (2012) In vitro transcription activities of Pol IV, Pol V, and RDR2 reveal coupling of Pol IV and RDR2 for dsRNA synthesis in plant RNA silencing. Mol Cell 48(5):811–818.

23. Sydow JF, et al. (2009) Structural basis of transcription: mismatch-specific fidelity mechanisms and paused RNA polymerase II with frayed RNA. Mol Cell 34(6):710–721.

24. Law JA, Vashisht AA, Wohlschlegel JA, & Jacobsen SE (2011) SHH1, a homeodomain protein required for DNA methylation, as well as RDR2, RDM4, and chromatin remodeling factors, associate with RNA polymerase IV. PLoS Genet 7(7):e1002195.

25. Goodman MF, Keener S, Guidotti S, & Branscomb EW (1983) On the enzymatic basis for mutagenesis by manganese. J Biol Chem 258(6):3469–3475.

26. El-Deiry WS, Downey KM, & So AG (1984) Molecular mechanisms of manganese mutagenesis. Proc Natl Acad Sci U S A 81(23):7378–7382.

27. Lou DI, et al. (2013) High-throughput DNA sequencing errors are reduced by orders of magnitude using circle sequencing. Proc Natl Acad Sci U S A 110(49): 19872–19877.

28. Acevedo A & Andino R (2014) Library preparation for highly accurate population sequencing of RNA viruses. Nat Protoc 9(7): 1760–1769.

29. Acevedo A, Brodsky L, & Andino R (2014) Mutational and fitness landscapes of an RNA virus revealed through population sequencing. Nature 505(7485):686–690.

30. Traverse CC & Ochman H (2016) Conserved rates and patterns of transcription errors across bacterial growth states and lifestyles. Proc Natl Acad Sci U S A 113(12):3311–3316.

31. de Mercoyrol L, Corda Y, Job C, & Job D (1992) Accuracy of wheat-germ RNA polymerase II. General enzymatic properties and effect of template conformational transition from right-handed B-DNA to left-handed Z-DNA. Eur J Biochem 206(1):49–58.

32. Bartel DP (2009) MicroRNAs: target recognition and regulatory functions. Cell 136(2):215–233.

33. Qi Y, et al. (2006) Distinct catalytic and non-catalytic roles of ARGONAUTE4 in RNA-directed DNA methylation. Nature 443(7114):1008–1012.

34. Wierzbicki AT, Haag JR, & Pikaard CS (2008) Noncoding transcription by RNA polymerase Pol IVb/Pol V mediates transcriptional silencing of overlapping and adjacent genes. Cell 135(4):635–648.

35. Wierzbicki AT, et al. (2012) Spatial and functional relationships among Pol V-associated loci, Pol IV-dependent siRNAs, and cytosine methylation in the Arabidopsis epigenome. Genes & development 26(16): 1825–1836.

36. Bohmdorfer G, et al. (2016) Long non-coding RNA produced by RNA polymerase V determines boundaries of heterochromatin. Elife 5.

37. Wendte JM & Pikaard CS (2016) Targeting Argonaute to chromatin. Genes Dev 30(24): 2649–2650.

38. Lahmy S, et al. (2016) Evidence for ARGONAUTE4-DNA interactions in RNA-directed DNA methylation in plants. Genes Dev 30(23):2565–2570.

39. Pikaard CS, Haag JR, Pontes OM, Blevins T, & Cocklin R (2013) A Transcription Fork Model for Pol IV and Pol V-dependent RNA-Directed DNA Methylation. Cold Spring Harb Symp Quant Biol.

40. Sousa R & Padilla R (1995) A mutant T7 RNA polymerase as a DNA polymerase. EMBO J 14(18):4609–4621.

41. Wei W, et al. (2012) A role for small RNAs in DNA double-strand break repair. Cell 149(1):101–112.

42. Walmacq C, et al. (2009) Rpb9 subunit controls transcription fidelity by delaying NTP sequestration in RNA polymerase II. J Biol Chem 284(29):19601–19612.

43. Kaplan CD, Larsson KM, & Kornberg RD (2008) The RNA polymerase II trigger loop functions in substrate selection and is directly targeted by alpha-amanitin. Mol Cell 30(5):547–556.

44. Bar-Nahum G, et al. (2005) A ratchet mechanism of transcription elongation and its control. Cell 120(2):183–193.

45. Zakharova N, Bass I, Arsenieva E, Nikiforov V, & Severinov K (1998) Mutations in and monoclonal antibody binding to evolutionary hypervariable region of Escherichia coli RNA polymerase beta' subunit inhibit transcript cleavage and transcript elongation. J Biol Chem 273(38):24912–24920.

46. Sievers F, et al. (2011) Fast, scalable generation of high-quality protein multiple sequence alignments using Clustal Omega. Mol Syst Biol 7:539.

47. Goujon M, et al. (2010) A new bioinformatics analysis tools framework at EMBL-EBI. Nucleic Acids Res 38(Web Server issue):W695–699.

48. Li H & Durbin R (2009) Fast and accurate short read alignment with Burrows-Wheeler transform. Bioinformatics 25(14):1754–1760.

